## Supplementary material for "The normal human lymph node cell classification and landscape defined by high-dimensional spatial proteomics": Spatial cell classification of the hLN-Supplementary Data

Affiliations:
1 Istituto di Bioimmagini e Sistemi Biologici Complessi (IBSBC) – CNR Via F.lli Cervi, 93 20090 Segrate (MI).
2 NBFC, National Biodiversity Future Center, Palermo 90133, Italy.
3 Laboratorio di Data Science and Bioinformatics, IRCCS Istituto delle Scienze Neurologiche di Bologna – AUSL BO Ospedale Bellaria, Via Altura 3 - 40139 Bologna - Italy.
4 Pathology Unit, Department of Molecular and Translational Medicine-DMMT, University of Brescia, via Branze, 43, 25123 Brescia, Italy.
5 Pathology Unit, ASST Spedali Civili Di Brescia, Brescia,Italy.
6 The Leuven Institute for Single-cell Omics (LISCO), KULeuven, Leuven, Belgium.
7 Translational Cell and Tissue Research Unit, Department of imaging and pathology, KULeuven, Leuven, Belgium.
8 Pathology Department, University Hospital of Leuven, Leuven, Belgium.
9 Department of Immunobiology, Yale University, New Haven, Connecticut, USA.
10Department of Laboratory Medicine, Yale University, New Haven, Connecticut, USA.
11 Columbia University Irving Medical Center and New York Presbyterian Hospital, NewYork, NY, USA.
12 Department of Mathematics and Applications, University of Milano Bicocca, 20126 Milan, Italy.
13 Department of Experimental Oncology, European Institute of Oncology IRCCS, 20139 Milan, Italy
14 Department of Experimental, Diagnostic and Specialty Medicine, University of Bologna, 40127 Bologna, Italy.
15 Pathology, Department of Medicine and Surgery, Universitá di Milano-Bicocca, 20900 Monza (MI), Italy.

† These authors contributed equally to this work
¶ present address: Pathology, Saint Louis University School of Medicine, Saint Louis, MO, USA

Document S1: Supplementary figures, tables and legends.

### Supplementary TABLE LEGENDS

Table S1
Clinicopathologic data of the patients and samples. The table contains an identifying UPN number, age, sex, ethnicity, site, LN diameter, clinicopathologic diagnosis in short, links to whole slides and TMA H&E images in an external site.

Table S2
Primary and secondary antibody table. The table lists antibody and target names, clone and species, manufacturer, lot, RRID number, fluorescence channel and round number, specification about distribution, subcellular location, links to HGNC protein and Human Protein Atlas.

Table S3
Cell classification of whole LN and TMA cores with BRAQUE$^{global}$ (BRAQUE1). The cell classification and number of informative cases are listed; each case, names with the UPN number, lists the total cell numbers, the percentage and the number of clusters belonging to that cell type. At the bottom, summary values for each case are provided.

Table S4
Criteria to group cell types classified by BRAQUE1 for BRAQUE2 analysis. Assignment of cell types to a broad cell type category. Each cell type identified in BRAQUE$^{global}$ is given a broad cell type name, so that BRAQUE$^{subclass}$ can be run of the clusters belonging to that broad cell type.

Table S5
Comprehensive granular classification of LN cells (BRAQUE1). Comprehensive catalog of cell types from BRAQUE$^{global}$ classification of 6 whole LN. Note that the cell types do differ from the final classification after BRAQUE subclassification.

Table S6
Antibody list and subpanel allocations. The table contains a list of primary antibodies, abbreviation, list of panels formation for cell type BRAQUE$^{subclass}$ analysis as well as targeted panels for cell type subclassification.

Table S7
Cluster classification: all clusters for whole LNs. Clusters number, cell content, cell classification, phenotype, spatial allocation for all WLN samples, with BRAQUE2 and BRAQUE1.

Table S8
Comprehensive catalog of cell types. This is an extended version of Table 1.
The table lists all cell types identified by BRAQUE$^{subclass}$ and for stromal cells BRAQUE$^{global}$.
For each cell type the following are listed:
- A name and a description
- The preferential location of the cell type, calculated on all cells of that type in all clusters. Note that an extended list of zones has been used.
- The markers frequency, divided into three tiers (≥70%, 50-69% and 30-49%), calculated on the frequency in all clusters of that cell type
- The list of markers ranking first to fifth, irrespectively of the number of clusters in which they rank on top.
- The total number of cells and the total number of clusters for that cell type
- A list of statistically significative neighboring cells, at 22 AND 44 pixels (10 AND 20 µm). “Self” indicate self-neighboring cells.

- A list of other cell populations which are statistically neighbors and have a statistically significant overlap with the cell type in question. “Self” indicate self-overlapping populations.
- UBERON: Uber-anatomy Ontology definition according to the Cell Ontology and Human Cell Atlas initiative (https://rdcu.be/dM6hW).
- Markers not listed are either not significant/negative or present in <30% of clusters. For single cluster information see Table S7.

Table S9
Neighborhood relationship between cell types. The table contains all statistical significance values for each cell type (rows) versus the others (columns). The table contains all values, however, only the values equal or smaller than p 8.094544277157197e-06 (approx. 0.0000081) are shown as a red box. NB: the table contains all the numbers, which are not visualized because of conditional formatting.

Table S10
Overlap between cell types. The table contains all statistical significance values for each cell type (rows) versus the others (columns). The table contains all values, however, only the values of significance “0” which have a neighborhood value equal or smaller than p 8.094544277157197e-06 (approx. 0.0000081) are shown as a green box. NB: the table contains all the numbers, which are not visualized because of conditional formatting.

Table S11
List of cases in which TCF7+ PAX5+ cells are found. List of samples containing TCF7+ PAX5+ cells by IF image analysis at low power.

Table S12
Spatial frequency allocation of cell types. Each cell type is distributed by frequency on the spatial landscape landmarks (extended list). Allocation for individual LN is also reported.

Table S13
Presence of Fairy Circles in whole LN sections and 2mm TMA cores, with clinicopathologic data. Whole sections and TMA cores are evaluated for presence and conformation of nodular paracortical aggregates (Fairy Circles) by examining multicolor IF images containing HLA-DR, CD1c and CD68 or DAPI. The UPN identifier, the type of pathology and the presence of FC is listed.

Table S14
Definitions and description of LN anatomical zones. Spatial definition and description of the anatomical structures to which the cell types are assigned.

Table S15
Comparison of cell segmentation with CyBorgh or CellPose2, followed by cell clustering by BRAQUE on three LN TMA cores. $BRAQUE^{global}$ was run on each of three TMA cores after segmentation with two different segmentation algorithms. For each case and each algorithm total numbers, percentage and number of clusters are provided for each cell type identified.

Table S16
Comparison of cell clustering by BRAQUE on a TMA core with noise. Comparison of the effect of AF subtraction on BRAQUE analysis. A LN TMA core was processed with $BRAQUE^{global}$

before and after AF was subtracted from the images. Total cell number, percentage and number of clusters are given for each cell type identified.

Table S17
Effect of Ab panel variations on the cell classification on two whole LN, UPN107 and UPN108. The table contains the composition of targeted panels for cell type subclassification, the results for two LN.

Table S18
Comprehensive granular classification of LN cells. Comprehensive catalog of cell types and marker distribution by frequency in total cells (not total clusters).
For each cell type the following are listed:
- A name and a description
- The preferential location of the cell type, calculated on all cells of that type in all clusters.
- The markers frequency, divided into three tiers (≥70%, 50-69% and 30-49%), calculated on the frequency in all cells of that cell type
- The list of markers ranking first to fifth, irrespectively of the number of clusters in which they rank on top.
- The total number of cells and the total number of clusters for that cell type

Table S19
Harmonized subsets by broad cell types. Comprehensive catalog of cell types, broadly defined. The table lists all broad cell types identified by BRAQUEglobal.
For each cell type the following are listed:
- A name and a description
- The markers frequency, divided into three tiers (≥70%, 50-69% and 30-49%), calculated on the frequency in all clusters of that cell type

Table S20
Spatial frequency allocation of cell types. Each cell type is distributed by frequency on the spatial landscape landmarks (extended LN zone list). Allocation for individual LN is also reported.

### Supplementary FIGURE LEGENDS

**Figure S1 (related to Figure 1)**
**A**: Low-power images of a whole LN (UPN106) stained for 81 markers. Each 8-bit image was enhanced via the Enhance-Contrast command in Fiji (0.35% saturated pixels). In each image, the biomarker, the fluorochrome and the round number is shown at the bottom. Note duplicate fluorochrome images for some markers. Scale bar 5 mm.
**B**: high-power magnification of 78 images from A, used for the analysis. In each image, the biomarker and the fluorochrome is shown at the bottom. Scale bar 500 µm.
**C**: Bivariate correlation matrix of all markers. Each marker is correlated against all others and the result plotted according to the scale on the right. On top and on the left the fluorochrome for imaging (blue: BV480; green: Alexa 488; orange: Rhodamine RedX; red: Alexa 647) and the staining round number are listed for each marker. Data is from UPN107. Image plot

representing the correlation matrix of all markers (Antibodies), one towards all, obtained with the "corplot" function in R (reordered according to AOE, the angular order of the eigenvectors). Divergent color palette shows in blue positive correlation, in red negative correlation. Dot size corresponds to the value of correlation. Notably, T-cell and B-cell markers clusterize separately in the table (top left corner).
**D**: Graphic representation of labeling across cell types for the panel of biomarkers used in this work. The colored circles represent CD4, CD8, B-cells (including plasma cells), innate immune cells (cDC, pDC, ILC3) and myelomonocytic cells. At the center, an inverted pentagram represents multiple lineage co-expression and is reported for clarity at the left of the Voronoi graph. Markers names are allocated according to the cell type distribution. The assignment is approximative. For additional information see Supplementary Table 2.

**Figure S2 (related to Figure 2)**
**A:** Flowchart for the assignment of the TCF7 level category. Non-proliferating CD4 or CD8 T-cells are classified based on high, low and intermediate levels of TCF7, as depicted by the cluster signal intensity profile (red), compared to the sample population (blue).
**B:** Flowchart for the assignment of the subset type for CD4 and CD8 T-cell clusters upon subclustering. The presence/absence of subclassifying markers is drawn from the BRAQUE statistical output (see[10, 78])

**Figure S3 (related to Figure 2)**
**A:** Spatial distribution of CD4 T-cell subsets on five whole LN. Each population, the sum of the individual clusters, is shown in red over a gray shadow representing all CD4 T-cells. Each dot is a magnified cell.
**B:** Spatial distribution of CD8 T-cell subsets on five whole LN. Each population, the sum of the individual clusters, is shown in red over a gray shadow representing all CD8 T-cells. Each dot is a magnified cell.

**Figure S4 (related to Figure 3 and 4)**
**A:** Spatial distribution of B-cell subsets on five whole LN. Each population, the sum of the individual clusters, is shown in red over a gray shadow representing all B-cells. Each dot is a magnified cell.
**B:** Spatial distribution of dendritic and non-myeloid innate immune cell subsets on six whole LN. Each population, the sum of the individual clusters, is shown in red over a gray shadow representing all dendritic cells. Each dot is a magnified cell.

**Figure S5 (related to Figure 3)**
**A:** A: Multiplex phenotype of CD5+ MZ B cells coexpressing PAX5 and TCF7. The top 16 images show serial stainings for the same area from the marginal follicular border of LN UPN104, showing a coordinated TCF7+ immature transitional B cell phenotype, highlighted by the circled cells. The bottom four serial images show a different area from the same LN close to a T-cell area, encompassing the border of a PAX5+ B cell follicle on the lower left and a T cell zone hosting numerous CD3+ T-cells coexpressing TCF7 and GATA3 (red circles), TCF7 but not GATA3 (green circles) and GATA3 but not TCF7 (blue circle). None of these T cells is PAX5+. The scale of each circle is 7 µm. The grayscale IF images have been inverted and adjusted for visibility.
**B:** Spatial allocation of CSR memory (CD27+), plasma cells (k + l) and all CD27neg B-cell subsets. CSR memory (CD27+; red), plasma cells (k + l; blue) and all CD27neg B-cell subsets (yellow) are allocated in the LN space. Single cells from each cell type represented as enlarged dots have been coalesced, the outline border darkened and superimposed in Adobe Photoshop. CSR memory B-cells and plasmacells show the least superimposition (dark violet), these latter

showing greater overlap with naive B-cells (green) at the mantle zone border. Note that all CD27+ B-cell subsets do not have neighborhood relationships with plasma cells at the single cell level.

**Figure S6 (related to Figure 3)**
**A:** Three TMA cores are shown as inverted IF image of nuclei (grey, DAPI), kappa light chain+ plasma cells (blue), lambda+ PC (red). The number of PC in each core is shown color-coded at the bottom of each core. Scale bar: 1mm.
**B:** A representative field from A is enlarged for each TMA core. Top row: intensity thresholded kappa (blue) or lambda (red) PC are superimposed onto the DAPI inverted image (grey). Middle row: raw IF images of the same field is shown as a RGB composite with nuclear DAPI (blue), kappa (red) and lambda (green) PC. The PC count with CyBorgh for UPN33_4 is 1473 (kappa) and 1268 (lambda), with CellPose 1362 (kappa) and 862 (lambda). PC count (CyBorgh) for UPN31_5 are 1048 (kappa) and 1541 (lambda). For UPN29_4 are 934 (kappa) and 2779 (lambda). Bottom row: a segmentation mask (CyBorgh) is superimposed to the middle row IF composite.
**C:** The double-scale graph shows the kappa/lambda ratio for ISH for light chain RNA (grey bars, scale on the left). The triangles show the percentage of ISH-positive signals over the whole LN area (scale on the right).
**D:** low-power images of the ISH for light chain RNA on serial LN sections.

**Figure S7 (related to Figure 5)**
**A:** Spatial distribution of myelomonocytic subsets on six whole LN. Each population, the sum of the individual clusters, is shown in red over a gray shadow representing all myelomonocytic cells. Each dot is a magnified cell
**B:** Spatial distribution of stromal cell subsets on six whole LN. Each population, the sum of the individual clusters, is shown in red over a gray shadow representing all stromal cells. Each dot is a magnified cell.

**Figure S8 (related to Figure 6)**
**A:** Neighborhood relationships at 22 and 44 pixels. A high magnification H&E image of a LN shows four index cells (in[superscript]) around whose center an inner 22 pixels circle and an outer 44 pixels circle are drawn. A red star marks neighboring nuclei encroached by the 22 pixel circle, a blue star the ones for the 44 pixel circle. Two macrophages (i1 and i3) contact one or no nuclei in the inner circle and 6 and 9 nuclei in the outer circle. A lymphocyte (i2) contacts five and nine nuclei in the inner and outer circle respectively. An endothelial cell (i4) contacts a single close neighboring nucleus and five nuclei in the outer circle. Scale: 0.45 µm per pixel.
**B:** Overlap scheme. By choosing an interacting area the overlap measure is the percentage of shared interacting areas among different cells. The overlap measure is bounded between 0% and 100%, and interactions were defined significantly only if 10 µm overlap (d1) AND 20 µm overlap (d2) resulted both significantly enhanced with respect to randomly placed cells (tested with bootstrap method). In this scheme, ***i*** is the index cell evaluated for overlap with two ***A*** cells belonging to a different cell type; those fulfill the statistical criteria for overlap, while the ***B*** cell does not.

**Figure S9 (related to Figure 6)**
Comparison of the significance of neighborhood relationships at 22 or 44 pixels for CD4+ T cells (**A**) or B cells (**B**). The figure contains all statistical significance values for each cell type (rows) versus the others (columns). Only the values equal or smaller than p 8.094544277157197e-06 (approx. 0.0000081) are shown. Color scale at the right. Statistically significant relationships at 22 pixels are in general captured and confirmed at 44 pixels as well, because of a larger capture

area. However, significant relationships at 44 pixels only (red squares) can be observed and represent spurious results from passers by cells. The neighborhood analysis for all cell types can be downloaded from UNIMIB Digital Commons repository Bicocca Open Archive Research Data (BOARD), doi: 10.17632/3ntbp3zdzh.1. 1 pixel = 0.45 µm.

**Figure S10 (related to Figure 6)**
**A:** Logarithm of odds ratio obtained from Fisher exact test. Only the values equal or smaller than p 8.094544277157197e-06 (approx. 0.0000081) are shown, where in order to combine p values from different thresholds and different lymph nodes the maximum p value was taken to show only most robust results. Color scale at the right.
**B:** Median distance of each cell type (rows) to the nearest cell of a given type (columns).To aggregate the results from different threshold and different lymph nodes the median of medians was taken. Color scale at the right.

**Figure S11 (related to Figure 4)**
Relationship between PNAd+ high endothelial venules (HEV) and Fairy Circles.
Three medium-power details from each of three lymph nodes are shown. The HEV are shown in white, the Fairy Circles are a commixtion of HLA-DR (red) and CD1c (green) positive cDC2 dendritic cells. Two small HLA-DR+ germinal centers are circled (dashed circle) in UPN105. Space bar : 200 µm.

**Figure S12 (related to Figure 1)**
Validation of the BRAQUE pipeline analysis.
**A:** Comparison of image segmentation with Cellpose 2.0 and CyBorgh. Three TMA cores (UPN26, 32 and 33) were segmented using Cellpose 2.0 or CyBorgh (part of the BRAQUE pipeline). BRAQUE was run on the .csv data files and the clusters were classified. The mean cell type percentages ± SD are shown. For an explanation of the cell types, see Table S5.
**B:** Spatial and phenotypic discrimination of juxtaposed cells by BRAQUE. Two examples of cells juxtaposed are shown both in the UMAP and in the real space for UPN107. In each panel the clusters of endothelium and stroma (i, gray background), Lyve1+ macrophages and Lyve1+ endothelium (ii, white background) are plotted in contrasting colors, respectively on the UMAP (left) and real space (right). Details are magnified as insets. The markers significance distribution of two representative clusters for each pair are shown below the UMAP maps. Below each real space, a RGB composite of DAPI (blue) and color-coded representative markers for each cell type (red, green) are shown.
**i:** Endothelium (red) and stromal cells (blue) clusters are plotted in the UMAP space and on the real tissue space. The dashed squares are enlarged as inserts. The marker expression ranking of two adjacent clusters shown in the insert, clusters 103 (stromal) and cluster 105 (endothelial) is shown, with relevant markers highlighted. The triple IF RGB color image shows nuclear DAPI (blue), endothelial vWF (red) and fibroblasts CD248 (green). Note the very close relationship of CD248 fibroblasts with vWF+ endothelium. Scale bar: 500 µm.
**ii:** Lyve1+ Macrophages (red) and Lyve1+ endothelium (blue) clusters are plotted in the UMAP space and on the real tissue space. The dashed squares are enlarged as inserts. The marker expression ranking of two representative clusters, Lyve1+ Macrophages and Lyve1+ endothelium is shown, with relevant markers highlighted. The triple IF RGB color image shows nuclear macrophage PU1 (blue), endothelial Lyve1 (red) and macrophage CD163 (green). Note Lyve1+ and Lyve1- CD163+ macrophages displaying nuclear PU1. Scale bar: 100 µm
**C:** Comparison of cell classification by BRAQUE on AF-subtracted and raw images. 75 images from a TMA core (UPN32) were either autofluorescence-subtracted (noAF) or left untreated (with AF), then both were independently analyzed with BRAQUE and the clusters obtained were classified. The percentage for each cell type is reported.

**D:**Technical artifacts and excluded cells. Clusters classified as technical artifacts (“junk”) are plotted on real space. The scale represents pixels (0.45 µm/pixel). Bottom left: the color-coded clusters plotted on the UMAP space of UPN107. Black dots represent clusters/cells which are allocated in the HDBSCAN -1 cluster. The dotted area is magnified on the right.
**E:** Effect of the antibody panel composition on the detection of cell types by BRAQUE. Two LN (UPN107 and UPN108), stained with 78 markers and segmented, were analyzed with four panels: *i*) BRAQUE with the full panel (78 Abs), *ii*) non redundant, mutually exclusive markers (33 Abs), *iii*) a combination of anti-transcription factor antibodies and 14 key diagnostic markers (TF & friends; 41 Abs) and *iv*) antibodies selected for highest signal-to-noise ratio and cell type identification power (Only cleans; 68 Abs). The percentage of broad cell types in each panel is shown ±SD. Note the larger SD and an increase in junk and unclear clusters with the smaller panels. For details of the panel’s composition see Table S6, Table S17. Table S17 contains individual cases results with a detailed cell list.

**Figure S13 (related to Figure 1)**
Cluster analysis (Seurat) of 37 normal lymph node Tissue Microarray cores.
**A:** 37 TMA cores from 18 subjects stained in multiplex are clustered with Seurat. Individual clusters are grouped by cell type classification. Relevant markers for each group are highlighted (red rectangle and marker name at the bottom). Cluster 45 to 51 have been removed because of insufficient cell numbers each. Unclassifiable clusters 36, 38, 42 and 43 have been removed.
**B:** Cell type composition of 37 TMA cores from 18 subjects are clustered with Seurat. Individual clusters are grouped by cell type classification and the percentage composition is plotted. Note that unclear clusters (199992 cells, 10.7% of the total) have been removed before calculating the percentage of each type. The cell type names are different from the analysis with BRAQUE and are assigned by manual scoring of the heatmaps.
**C:** Comparison of global BRAQUE analysis and Seurat. 37 TMA cores from 18 subjects stained in multiplex are clustered with Seurat and BRAQUE. The cell types from BRAQUE are grouped in broad cell types for the comparison.

**Figure S14 (related to Figure 1)**
Cell type frequency in whole sections and TMA cores identified in 27 LN samples (6 whole LN and 21 2mm TMA cores) after $BRAQUE^{global}$ analysis on all cells in each sample, expressed as percentage ± SD. The cell types are separated into **A**: frequent cell types (i.e. ≥ 20 total clusters) and B: infrequent cell types (i.e. <20 total clusters). The numbers next to the bars are the number of informative cases for that cell type. Note frequency discrepancies, most notable in B, referable to the TMA area sampling selection. See also Table S3.

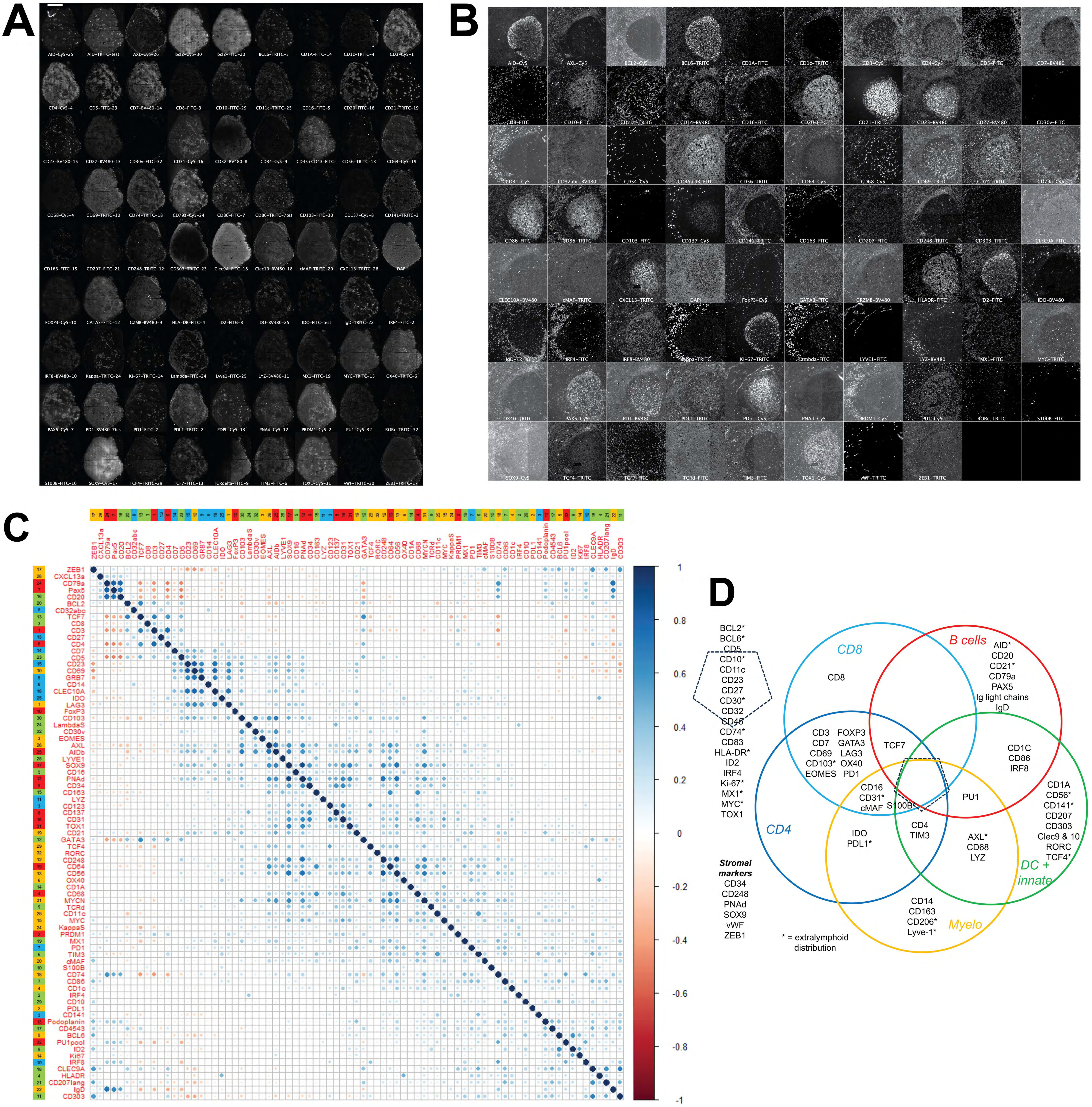
A
B
C
D
BCL2*
BCL6*
CD5
CD10*
CD11c
CD23
CD27
CD30*
CD32
CD45
CD74*
CD83
HLA-DR*
ID2
IRF4
Ki-67*
MX1*
MYC*
TOX1
Stromal markers
CD34
CD248
PNAd
SOX9
vWF
ZEB1
* = extralymphoid distribution
CD8
CD8
B cells
AID*
CD20
CD21*
CD79a
PAX5
Ig light chains
IgD
CD3
CD7
CD69
CD103*
EOMES
FOXP3
GATA3
LAG3
OX40
PD1
TCF7
CD1C
CD86
IRF8
CD16
CD31*
cMAF
S100B
PU1
CD1A
CD56*
CD141*
CD207
CD303
Clec9 & 10
RORC
TCF4*
CD4
IDO
PDL1*
CD4
TIM3
AXL*
CD68
LYZ
DC + innate
CD14
CD163
CD206*
Lyve-1*
Myelo

Figure S1

Figure S1 (related to Figure 1)
**A:** Low-power images of a whole LN (UPN106) stained for 81 markers. Each 8-bit image was enhanced via the Enhance-Contrast command in Fiji (0.35% saturated pixels). In each image, the biomarker, the fluorochrome and the round number is shown at the bottom. Note duplicate fluorochrome images for some markers. Scale bar 5 mm.
**B:** high-power magnification of 78 images from A, used for the analysis. In each image, the biomarker and the fluorochrome is shown at the bottom. Scale bar 500 μm.
**C:** Bivariate correlation matrix of all markers. Each marker is correlated against all others and the result plotted according to the scale on the right. On top and on the left the fluorochrome for
imaging (blue: BV480; green: Alexa 488; orange: Rhodamine RedX; red: Alexa 647) and the staining round number are listed for each marker. Data is from UPN107. Image plot representing the correlation matrix of all markers (Antibodies), one towards all, obtained with the “corplot” function in R (reordered according to AOE, the angular order of the eigenvectors). UPN107 dataset has been chosen as reference.Divergent color palette shows in blue positive correlation, in red negative correlation.Dot size corresponds to the value of correlation. The coloured bars (top and left) represent the IF channel of acquisition of each marker, (blue: BV480; green: Alexa 488; orange: Rhodamine RedX; red: Alexa 647) . Numbers inside bars indicate the round of staining with MILAN. Notably, T cell and B cell markers clusterize separately in the table (top left corner).
**D:** Graphic representation of labeling across cell types for the panel of biomarkers used in this work. The colored circles represent CD4, CD8, B cells (including plasma cells), innate immune cells (cDC, pDC, ILC3) and myelomonocytic cells. At the center, an inverted pentagram represents multiple lineage co-expression and is reported for clarity at the left of the Voronoi graph. Markers names are allocated according to the cell type distribution. The assignment is approximative. For additional information see Supplementary Table 2.

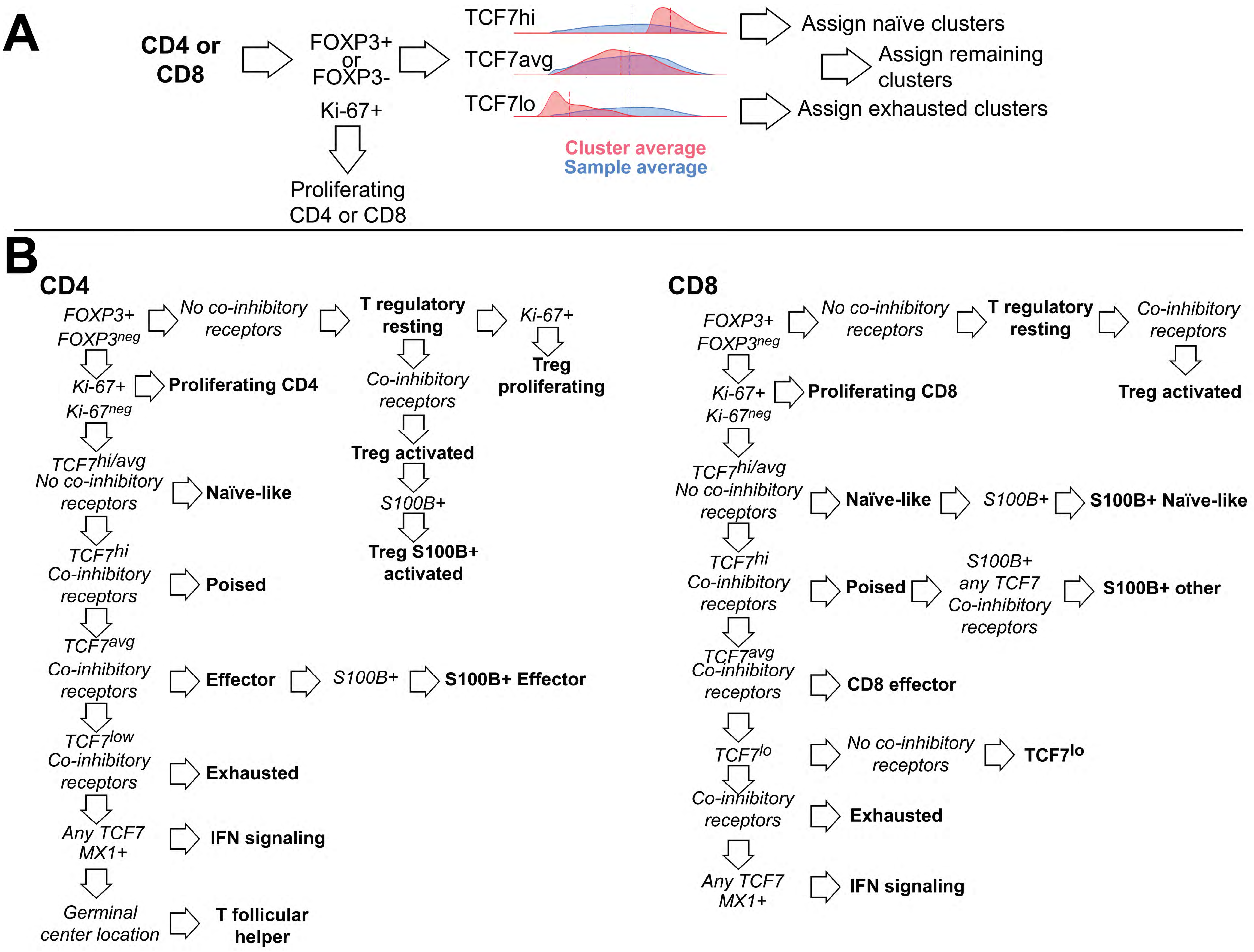


Figure S2 (related to Figure 2)

**A:** Flowchart for the assignment of the TCF7 level category. Non-proliferating CD4 or CD8 T-cells are classified based on high, low and intermediate levels of TCF7, as depicted by the cluster signal intensity profile (red), compared to the sample population (blue).

**B**: Flowchart for the assignment of the subset type for CD4 and CD8 T-cell clusters upon subclustering. The presence/absence of subclassifying markers is drawn from the BRAQUE statistical output.

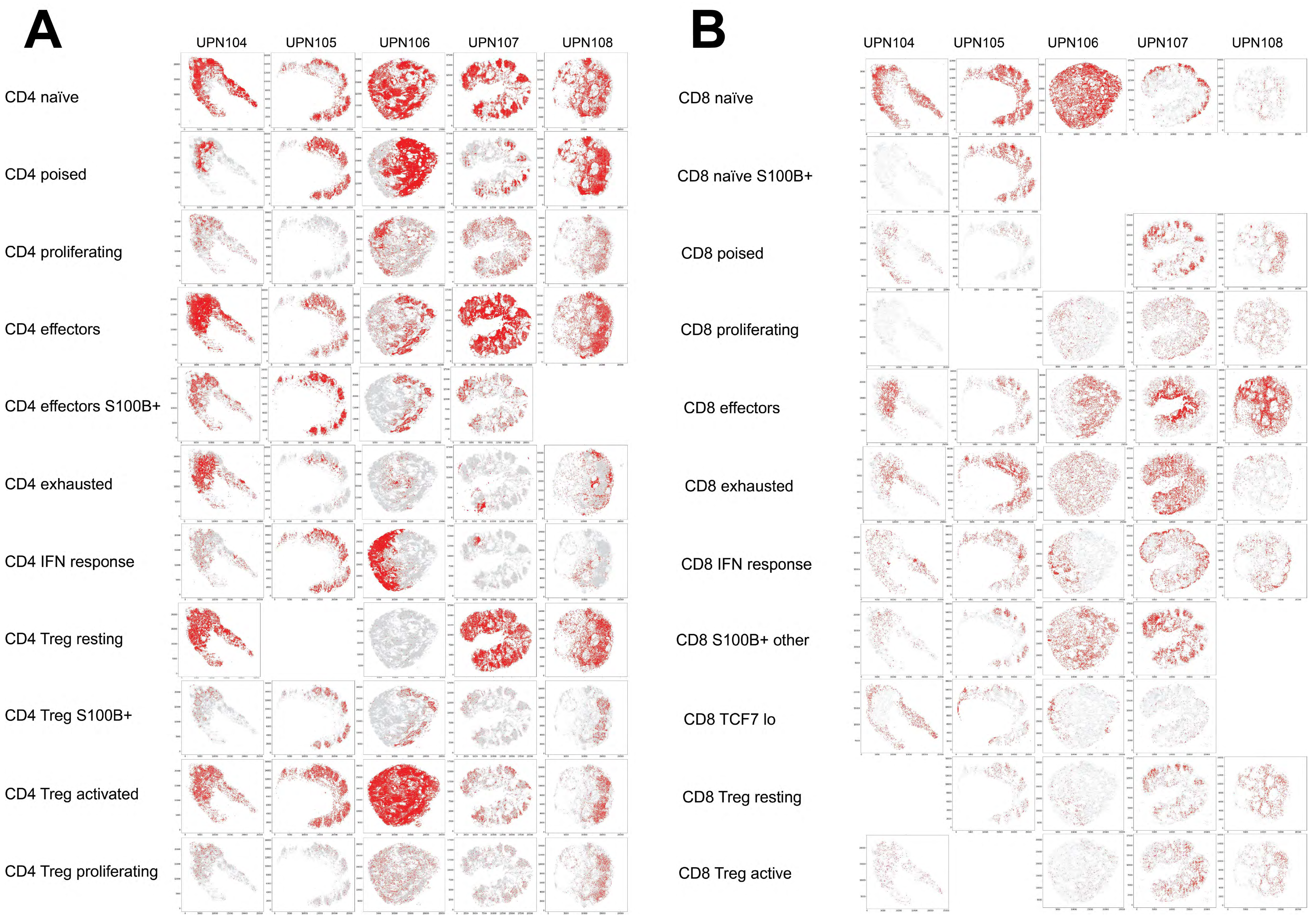


### Figure S3 (related to Figure 2)

**A:** Spatial distribution of CD4 T-cell subsets on five whole LN. Each population, the sum of the individual clusters, is shown in red over a gray shadow representing all CD4 T cells. Each dot is a magnified cell.

**B:** Spatial distribution of CD8 T-cell subsets on five whole LN. Each population, the sum of the individual clusters, is shown in red over a gray shadow representing all CD8 T cells. Each dot is a magnified cell.

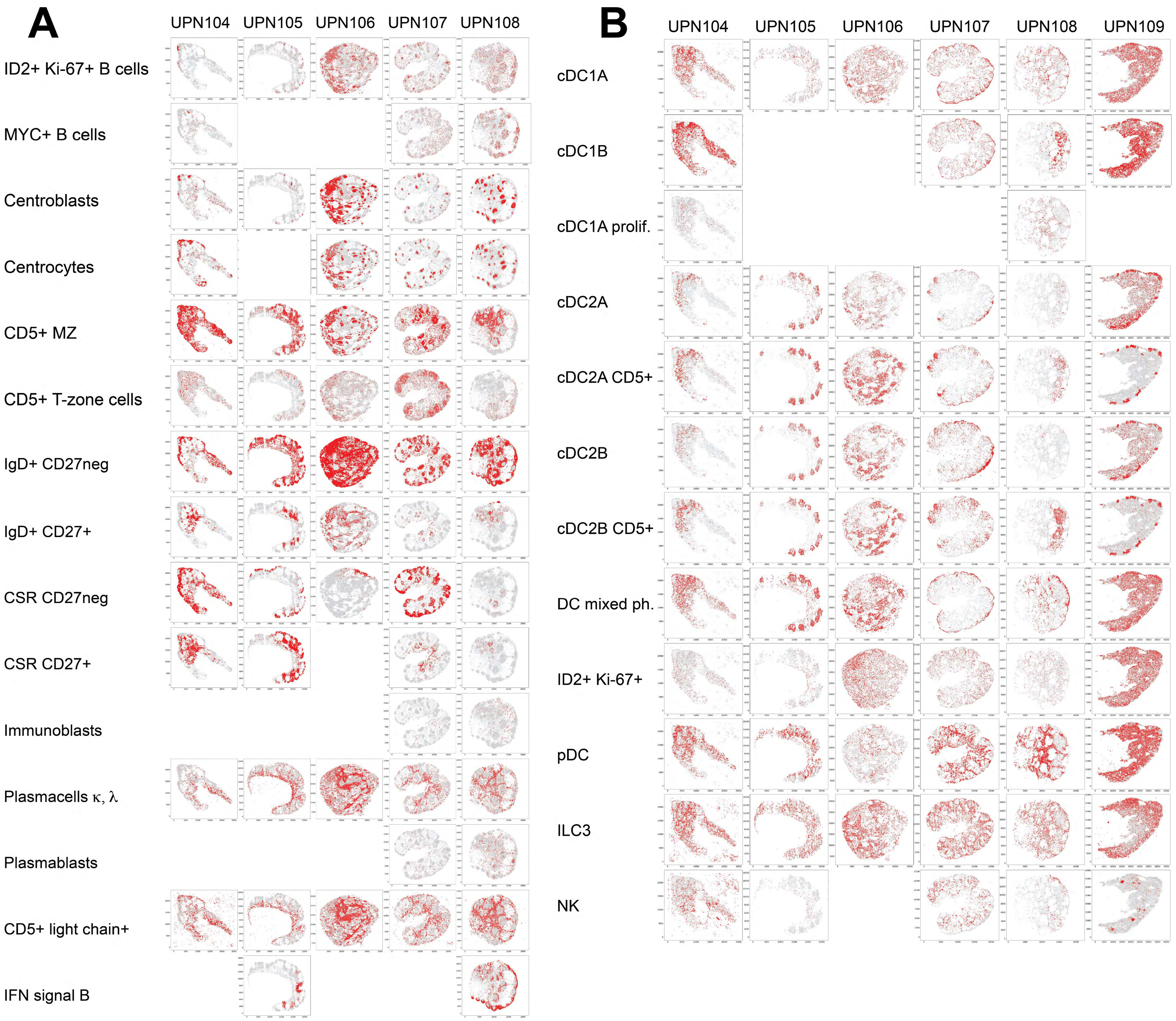


### Figure S4 (related to Figure 3 and 4)

**A:** Spatial distribution of B-cell subsets on five whole LN. Each population, the sum of the individual clusters, is shown in red over a gray shadow representing all B cells. Each dot is a magnified cell.

**B:** Spatial distribution of dendritic and non-myeloid innate immune cell subsets on six whole LN. Each population, the sum of the individual clusters, is shown in red over a gray shadow representing all dendritic cells. Each dot is a magnified cell.

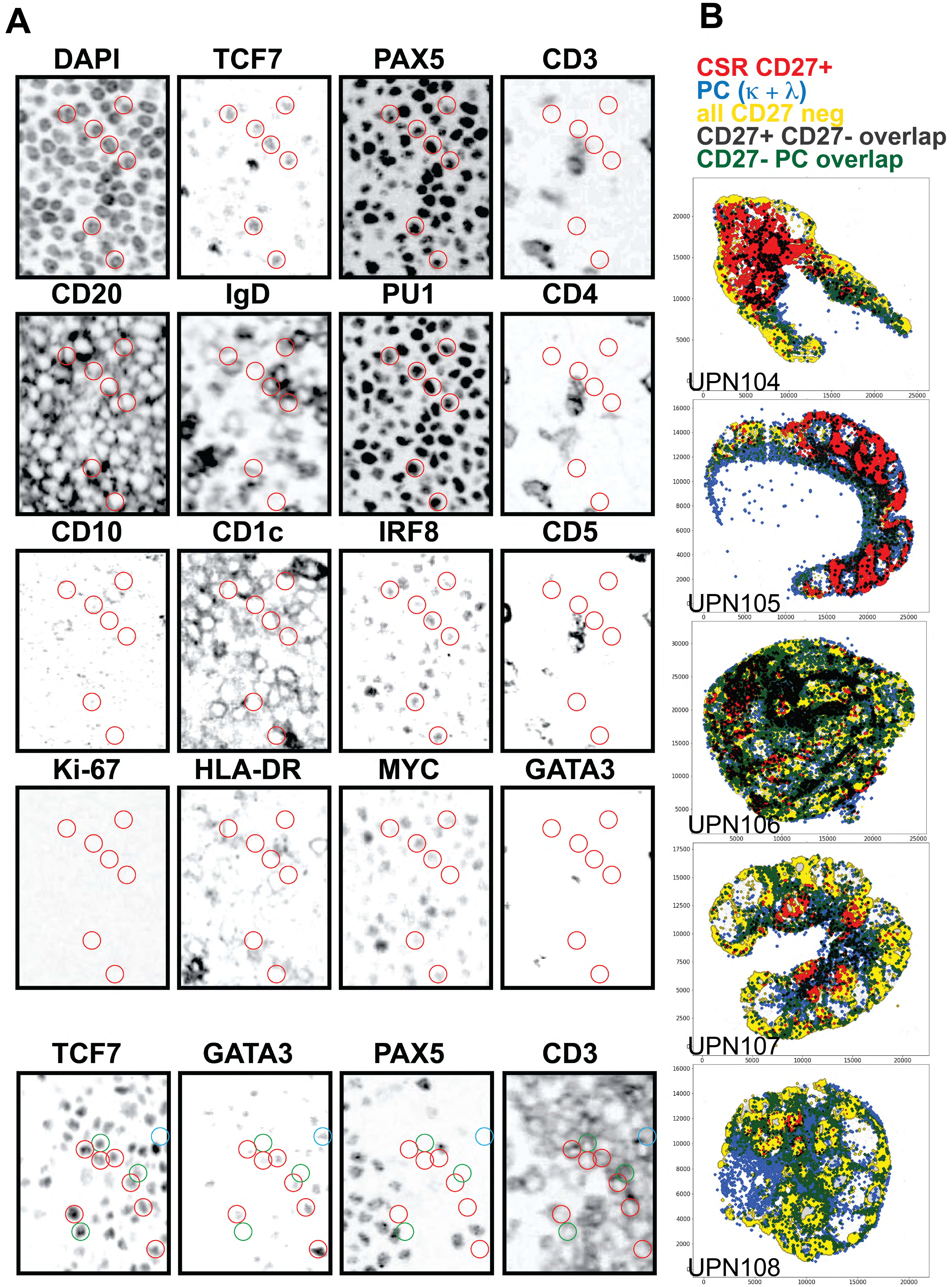
A
DAPI
TCF7
PAX5
CD3
CD20
IgD
PU1
CD4
CD10
CD1c
IRF8
CD5
Ki-67
HLA-DR
MYC
GATA3
TCF7
GATA3
PAX5
CD3
B
CSR CD27+
PC (κ + λ)
all CD27 neg
CD27+ CD27- overlap
CD27- PC overlap
UPN104
UPN105
UPN106
UPN107
UPN108

Figure S5

### Figure S5 (related to Figure 3)

**A:** Multiplex phenotype of CD5+ MZ B cells coexpressing PAX5 and TCF7. The top 16 images show serial stainings for the same area from the marginal follicular border of LN UPN104, showing a coordinated TCF7+ immature transitional B cell phenotype, highlighted by the circled cells. The bottom four serial images show a different area from the same LN close to a T-cell area, encompassing the border of a PAX5+ B cell follicle on the lower left and a T cell zone hosting numerous CD3+ T cells coexpressing TCF7 and GATA3 (red circles), TCF7 but not GATA3 (green circles) and GATA3 but not TCF7 (blue circle). None of these T cells is PAX5+. The scale of each circle is 7 µm. The grayscale IF images have been inverted and adjusted for visibility.

**B:** Spatial allocation of CSR memory (CD27+), plasma cells (k + l) and all CD27neg B cell subsets.

CSR memory (CD27+; red), plasma cells (k + l; blue) and all CD27neg B cell subsets (yellow) are allocated in the LN space. Single cells from each cell type represented as enlarged dots have been coalesced, the outline border darkened and superimposed in Adobe Photoshop. CSR memory B cells and plasmacells show the least superimposition (dark violet), while showing greater overlap with naive B cells (green) at the mantle zone border. Note that all CD27+ B cell subsets do not have neighborhood relationships with plasma cells at the single cell level.

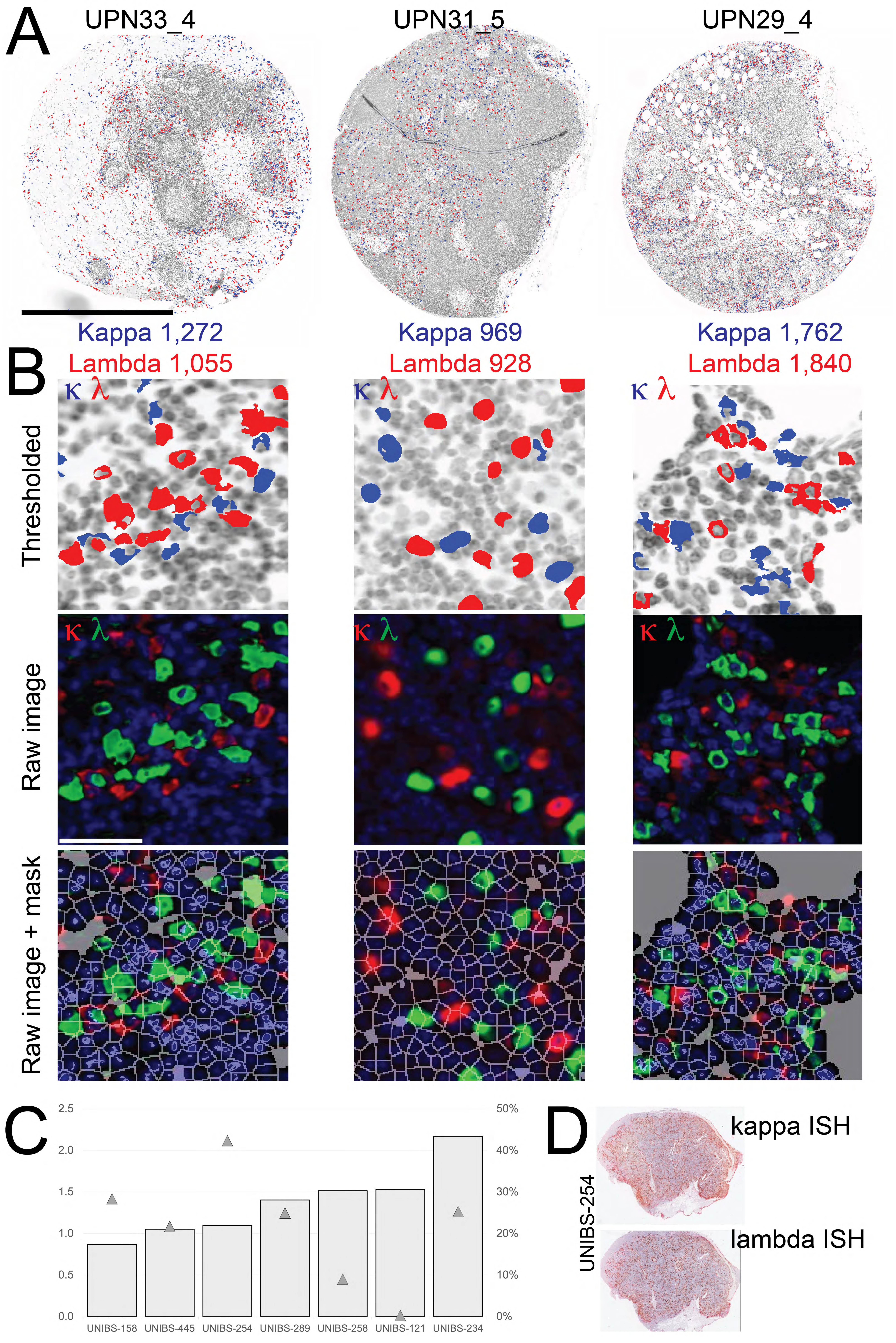
A
UPN33_4
UPN31_5
UPN29_4
Kappa 1,272
Lambda 1,055
Kappa 969
Lambda 928
Kappa 1,762
Lambda 1,840
B
Thresholded
Raw image
Raw image + mask
κ λ
C
2.5
2.0
1.5
1.0
0.5
0.0
50%
40%
30%
20%
10%
0%
UNIBS-158
UNIBS-445
UNIBS-254
UNIBS-289
UNIBS-258
UNIBS-121
UNIBS-234
D
UNIBS-254
kappa ISH
lambda ISH

Figure S6

### Figure S6 (related to Figure 3)

**A:** Three TMA cores are shown as inverted IF image of nuclei (grey, DAPI), kappa light chain+ plasma cells (blue), lambda+ PC (red). The number of PC in each core is shown color-coded at the bottom of each core. Scale bar: 1mm.
**B:** A representative field from A is enlarged for each TMA core. Top row: intensity thresholded kappa (blue) or lambda (red) PC are superimposed onto the DAPI inverted image (grey). Middle row: raw IF images of the same field is shown as a RGB composite with nuclear DAPI (blue), kappa (red) and lambda (green) PC. The PC count with CyBorgh for UPN33_4 is 1473 (kappa) and 1268 (lambda), with CellPose 1362 (kappa) and 862 (lambda). PC count (CyBorgh) for UPN31_5 are 1048 (kappa) and 1541 (lambda). For UPN29_4 are 934 (kappa) and 2779 (lambda). Bottom row: a segmentation mask (CyBorgh) is superimposed to the middle row IF composite.
**C:** The double-scale graph shows the kappa/lambda ratio for ISH for light chain RNA (grey bars, scale on the left). The triangles show the percentage of ISH-positive signals over the whole LN area (scale on the right).
**D:** low-power images of the ISH for light chain RNA on serial LN sections..

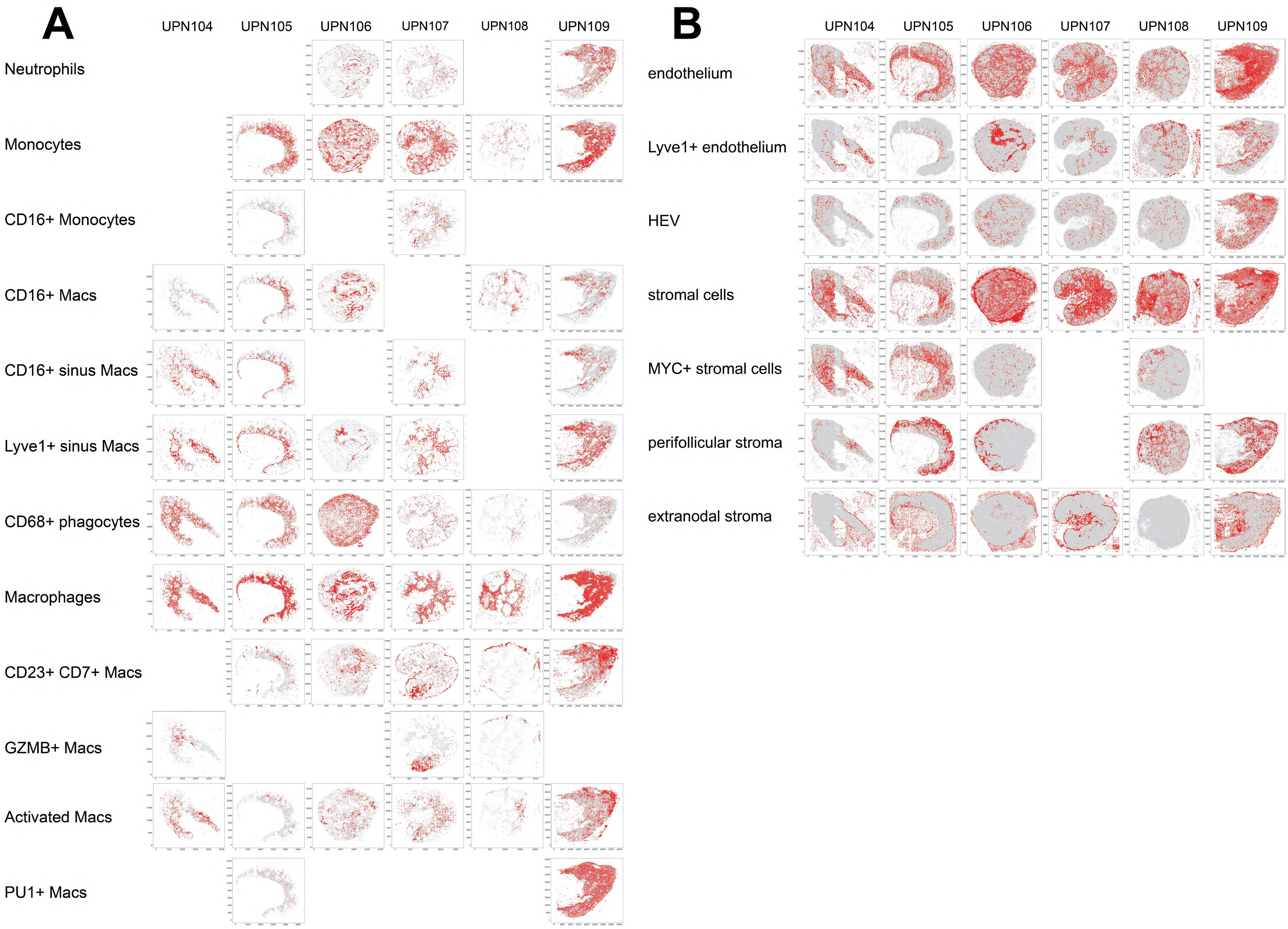


### Figure S7 (related to Figure 5)

**A:** Spatial distribution of myelomonocytic subsets on six whole LN. Each population, the sum of the individual clusters, is shown in red over a gray shadow representing all myelomonocytic cells. Each dot is a magnified cell

**B:** Spatial distribution of stromal cell subsets on six whole LN. Each population, the sum of the individual clusters, is shown in red over a gray shadow representing all stromal cells. Each dot is a magnified cell.

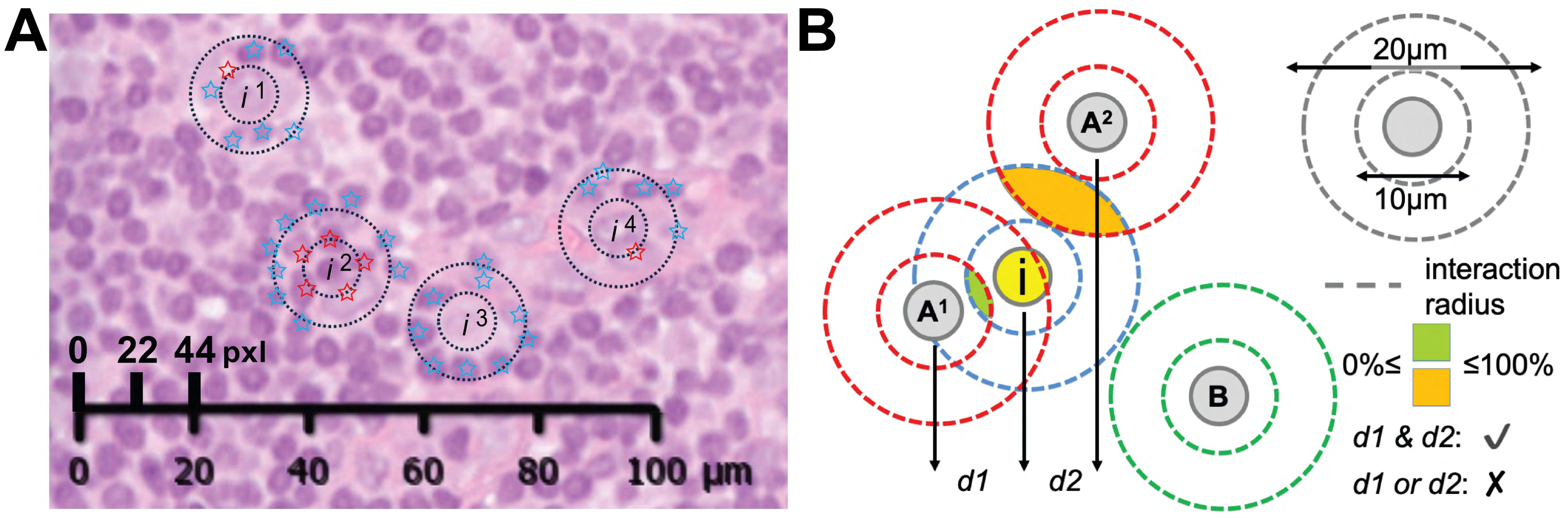


Figure S8 (related to Figure 6)

Neighborhood and cell type overlap (related to Figure 6)

**A:** Neighborhood relationships at 22 and 44 pixels. A high magnification H&E image of a LN shows four index cells (in[superscript]) around whose center an inner 22 pixels circle and an outer 44 pixels circle are drawn. A red star marks neighboring nuclei encroached by the 22 pixel circle, a blue star the ones for the 44 pixel circle. Two macrophages ($i^1$ and $i^3$) contact one or no nuclei in the inner circle and 6 and 9 nuclei in the outer circle. A lymphocyte ($i^2$) contacts five and nine nuclei in the inner and outer circle respectively. An endothelial cell ($i^4$) contacts a single close neighboring nucleus and five nuclei in the outer circle. Scale: 0.45 µm per pixel.

**B:** Overlap scheme. By choosing an interacting area the overlap measure is the percentage of shared interacting areas among different cells. The overlap measure is bounded between 0% and 100%, and interactions were defined significantly only if 10 µm overlap (d1) AND 20 µm overlap (d2) resulted both significantly enhanced with respect to randomly placed cells (tested with bootstrap method). In this scheme, i is the index cell evaluated for overlap with two A cells belonging to a different cell type; those fulfill the statistical criteria for overlap, while the B cell does not.

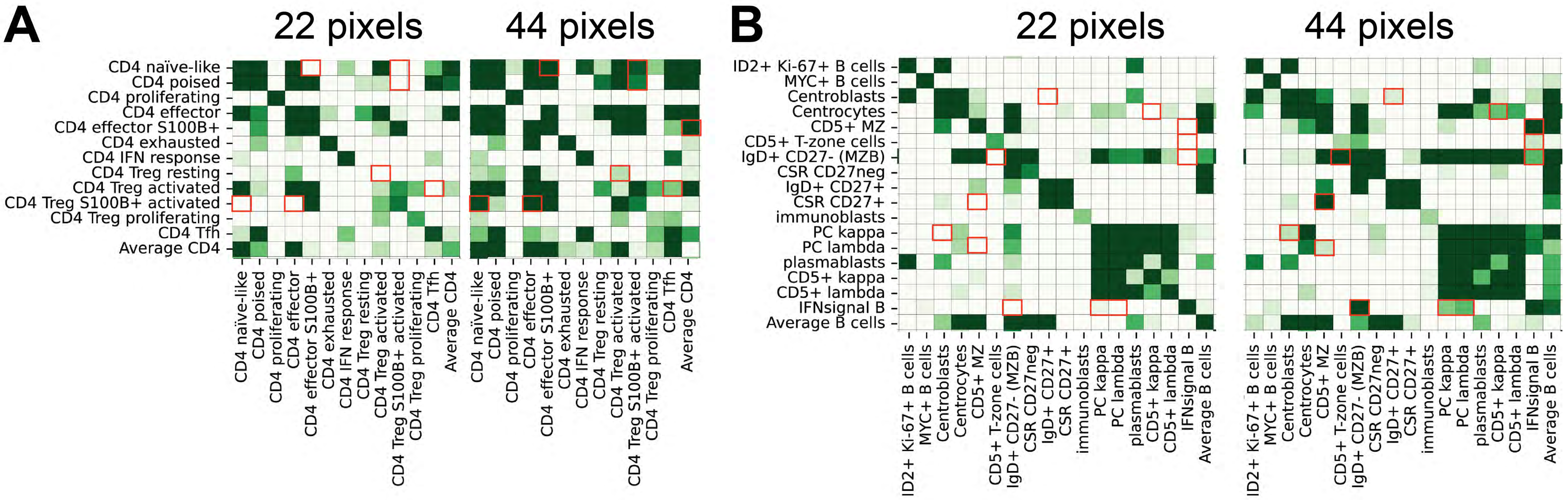


### Figure S9 (related to Figure 6)

Comparison of the significance of neighborhood relationships at 22 or 44 pixels for CD4+ T cells (**A**) or B cells (**B**). The figure contains all statistical significance values for each cell type (rows) versus the others (columns). Only the values equal or smaller than p 8.094544277157197e-06 (approx. 0.0000081) are shown. Color scale at the right. Statistically significant relationships at 22 pixels are in general captured and confirmed at 44 pixels as well, because of a larger capture area. However, significant relationships at 44 pixels only (red squares) can be observed and represent spurious results from passer-by cells. The neighborhood analysis for all cell types can be downloaded from UNIMIB Digital Commons repository Bicocca Open Archive Research Data (BOARD), doi: 10.17632/3ntbp3zdzh.1. 1 pixel = 0.45 μm.

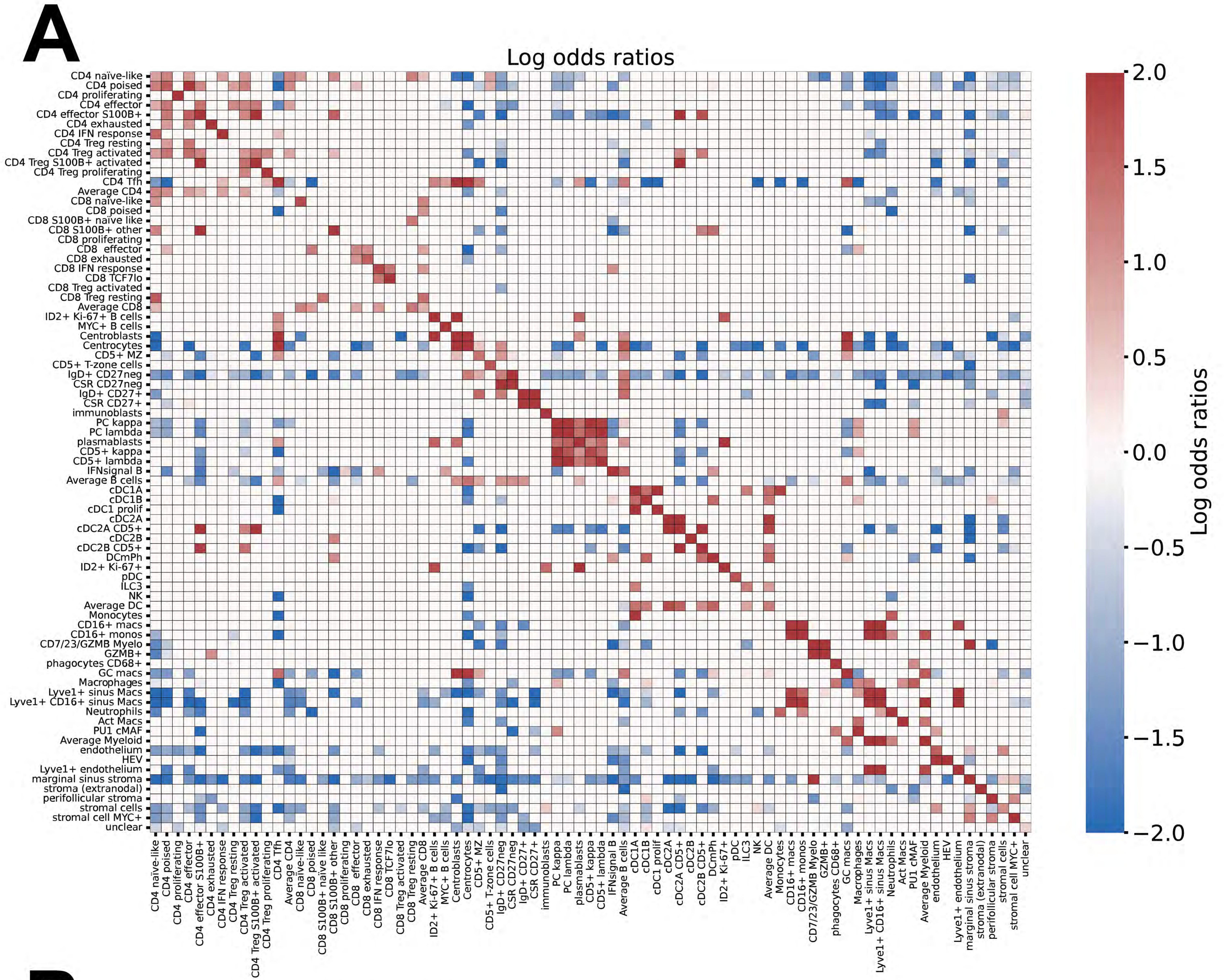
A
Log odds ratios
CD4 naïve-like
CD4 poised
CD4 proliferating
CD4 effector
CD4 effector S100B+
CD4 exhausted
CD4 IFN response
CD4 Treg resting
CD4 Treg activated
CD4 Treg S100B+ activated
CD4 Treg proliferating
CD4 Tfh
Average CD4
CD8 naïve-like
CD8 poised
CD8 S100B+ naïve like
CD8 S100B+ other
CD8 proliferating
CD8 effector
CD8 exhausted
CD8 IFN response
CD8 TCF7lo
CD8 Treg activated
CD8 Treg resting
Average CD8
ID2+ Ki-67+ B cells
MYC+ B cells
Centroblasts
Centrocytes
CD5+ MZ
CD5+ T-zone cells
IgD+ CD27neg
CSR CD27neg
IgD+ CD27+
CSR CD27+
immunoblasts
PC kappa
PC lambda
plasmablasts
CD5+ kappa
CD5+ lambda
IFNsignal B
Average B cells
cDC1A
cDC1B
cDC1 prolif
cDC2A
cDC2A CD5+
cDC2B
cDC2B CD5+
DCmPh
ID2+ Ki-67+
pDC
ILC3
NK
Average DC
Monocytes
CD16+ macs
CD16+ monos
CD7/23/GZMB Myelo
GZMB+
phagocytes CD68+
GC macs
Macrophages
Lyve1+ sinus Macs
Lyve1+ CD16+ sinus Macs
Neutrophils
Act Macs
PU1 cMAF
Average Myeloid
endothelium
HEV
Lyve1+ endothelium
marginal sinus stroma
stroma (extranodal)
perifollicular stroma
stromal cells
stromal cell MYC+
unclear
2.0
1.5
1.0
0.5
0.0
−0.5
−1.0
−1.5
−2.0
Log odds ratios

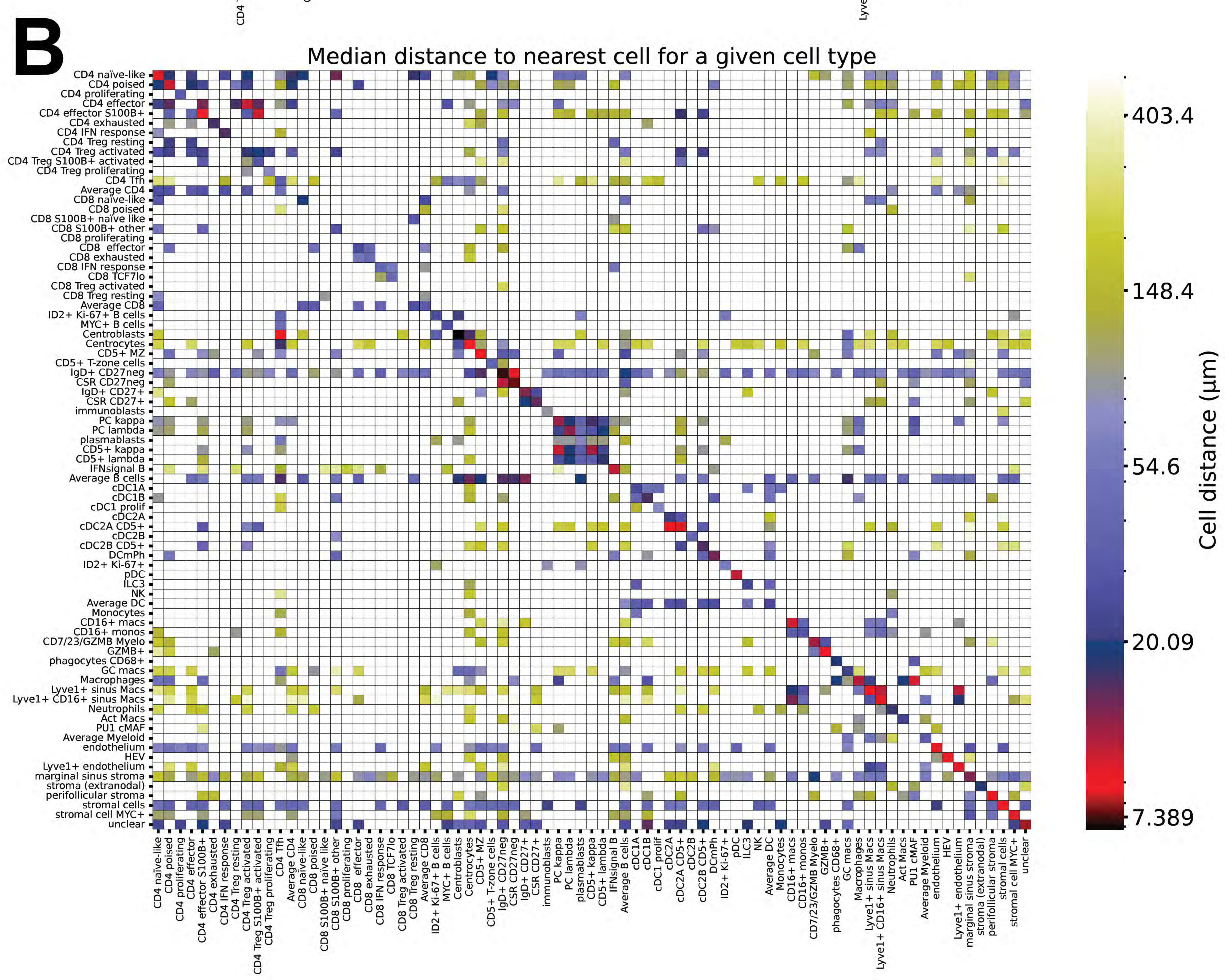
B
Median distance to nearest cell for a given cell type
CD4 naïve-like
CD4 poised
CD4 proliferating
CD4 effector
CD4 effector S100B+
CD4 exhausted
CD4 IFN response
CD4 Treg resting
CD4 Treg activated
CD4 Treg S100B+ activated
CD4 Treg proliferating
CD4 Tfh
Average CD4
CD8 naïve-like
CD8 poised
CD8 S100B+ naïve like
CD8 S100B+ other
CD8 proliferating
CD8 effector
CD8 exhausted
CD8 IFN response
CD8 TCF7lo
CD8 Treg activated
CD8 Treg resting
Average CD8
ID2+ Ki-67+ B cells
MYC+ B cells
Centroblasts
Centrocytes
CD5+ MZ
CD5+ T-zone cells
IgD+ CD27neg
CSR CD27neg
IgD+ CD27+
CSR CD27+
immunoblasts
PC kappa
PC lambda
plasmablasts
CD5+ kappa
CD5+ lambda
IFNsignal B
Average B cells
cDC1A
cDC1B
cDC1 prolif
cDC2A
cDC2A CD5+
cDC2B
cDC2B CD5+
DCmPh
ID2+ Ki-67+
pDC
ILC3
NK
Average DC
Monocytes
CD16+ macs
CD16+ monos
CD7/23/GZMB Myelo
GZMB+
phagocytes CD68+
GC macs
Macrophages
Lyve1+ sinus Macs
Lyve1+ CD16+ sinus Macs
Neutrophils
Act Macs
PU1 cMAF
Average Myeloid
endothelium
HEV
Lyve1+ endothelium
marginal sinus stroma
stroma (extranodal)
perifollicular stroma
stromal cells
stromal cell MYC+
unclear
403.4
148.4
54.6
20.09
7.389
Cell distance (μm)

Figure S10

### Figure S10 (related to Figure 6)

**A:** Logarithm of odds ratio obtained from Fisher exact test. Only the values equal or smaller than p 8.094544277157197e-06 (approx. 0.0000081) are shown, where in order to combine p values from different thresholds and different lymph nodes the maximum p value was taken to show only most robust results. Color scale at the right.
**B:** Median distance of each cell type (rows) to the nearest cell of a given type (columns).To aggregate the results from different threshold and different lymphnodes the median of medians was taken. Color scale at the right.

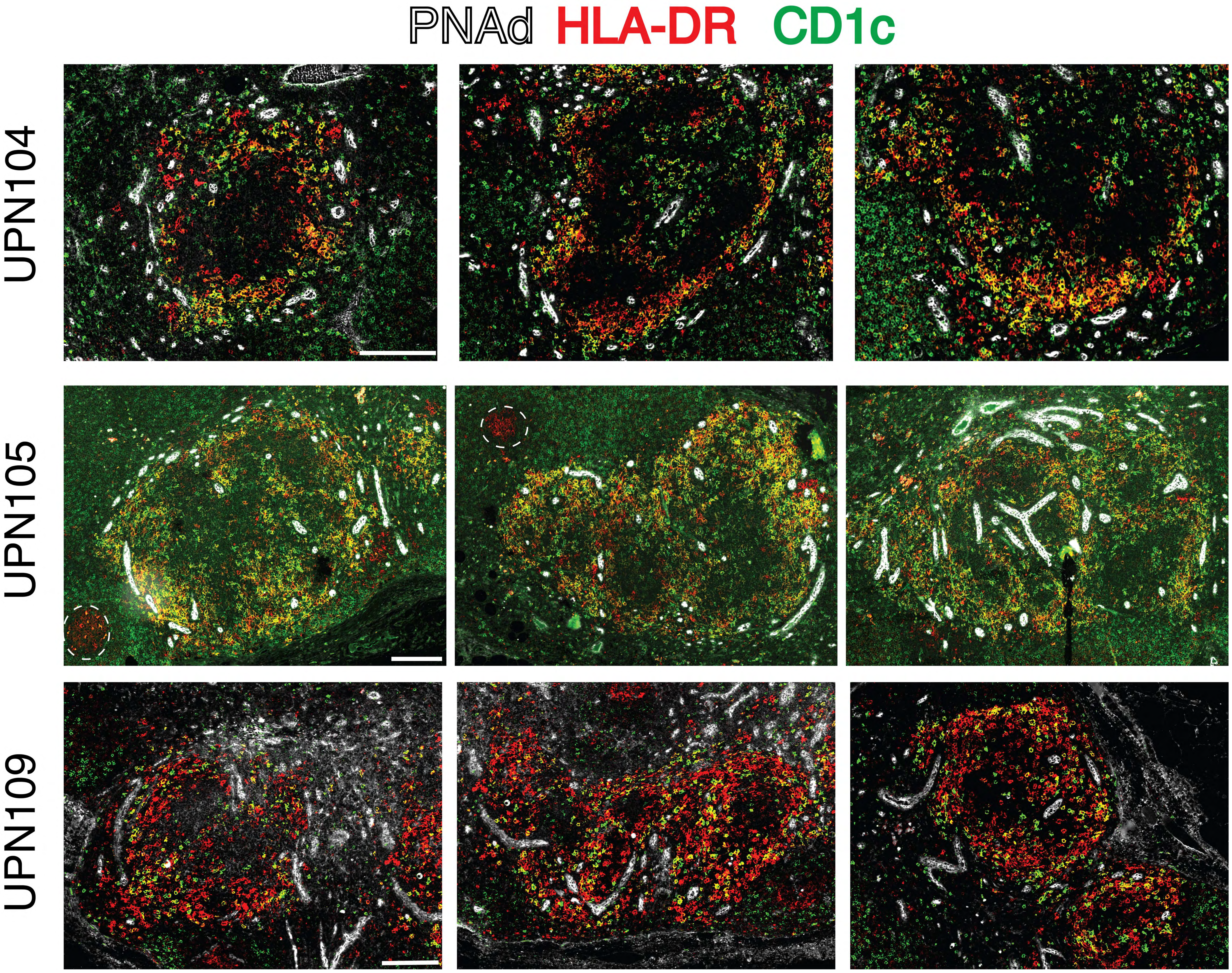


Figure S11 (related to Figure 4)
Relationship between PNAd+ high endothelial venules (HEV) and Fairy Circles.
Three medium-power details from each of three lymph nodes are shown. The HEV are shown in white, the Fairy Circles are a commixtion of HLA-DR (red) and CD1c (green) positive cDC2 dendritic cells. Two small HLA-DR+ germinal centers are circled (dashed circle) in UPN105.
Space bar : 200 μm.

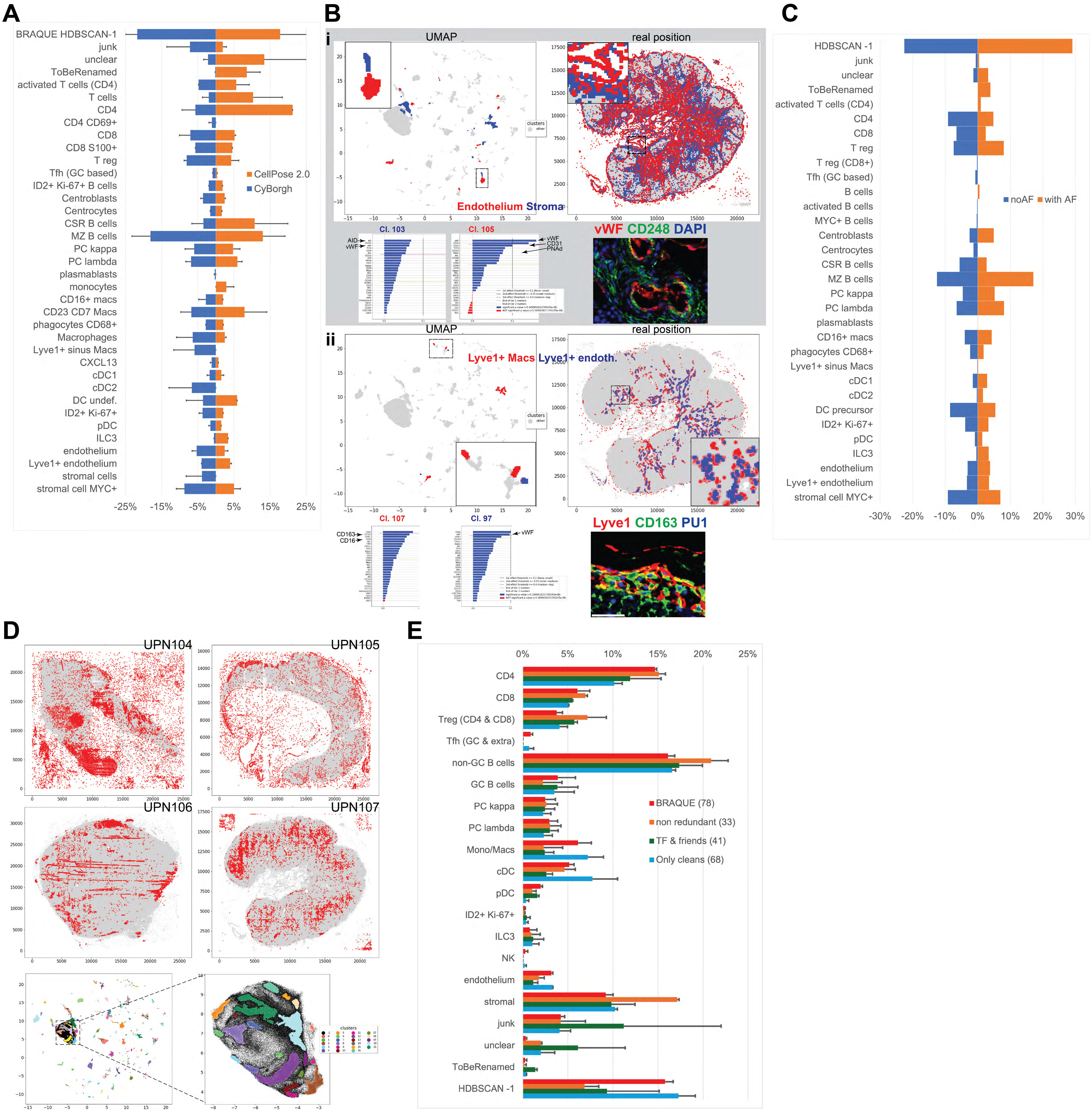
A
BRAQUE HDBSCAN-1
junk
unclear
ToBeRenamed
activated T cells (CD4)
T cells
CD4
CD4 CD69+
CD8
CD8 S100+
T reg
Tfh (GC based)
ID2+ Ki-67+ B cells
Centroblasts
Centrocytes
CSR B cells
MZ B cells
PC kappa
PC lambda
plasmablasts
monocytes
CD16+ macs
CD23 CD7 Macs
phagocytes CD68+
Macrophages
Lyve1+ sinus Macs
CXCL13
cDC1
cDC2
DC undef.
ID2+ Ki-67+
pDC
ILC3
endothelium
Lyve1+ endothelium
stromal cells
stromal cell MYC+
CellPose 2.0
CyBorgh
-25% -15% -5% 5% 15% 25%
B
i
UMAP
real position
Endothelium Stroma
Cl. 103
Cl. 105
AID
vWF
CD31
PNAd
vWF CD248 DAPI
ii
Lyve1+ Macs Lyve1+ endoth.
Cl. 107
Cl. 97
CD163
CD16
Lyve1 CD163 PU1
C
HDBSCAN -1
junk
unclear
ToBeRenamed
activated T cells (CD4)
CD4
CD8
T reg
T reg (CD8+)
Tfh (GC based)
B cells
activated B cells
MYC+ B cells
Centroblasts
Centrocytes
CSR B cells
MZ B cells
PC kappa
PC lambda
plasmablasts
CD16+ macs
phagocytes CD68+
Lyve1+ sinus Macs
cDC1
cDC2
DC precursor
ID2+ Ki-67+
pDC
ILC3
endothelium
Lyve1+ endothelium
stromal cell MYC+
noAF
with AF
-30% -20% -10% 0% 10% 20% 30%
D
UPN104
UPN105
UPN106
UPN107
clusters
E
0% 5% 10% 15% 20% 25%
CD4
CD8
Treg (CD4 & CD8)
Tfh (GC & extra)
non-GC B cells
GC B cells
PC kappa
PC lambda
Mono/Macs
cDC
pDC
ID2+ Ki-67+
ILC3
NK
endothelium
stromal
junk
unclear
ToBeRenamed
HDBSCAN -1
BRAQUE (78)
non redundant (33)
TF & friends (41)
Only cleans (68)

### Figure S12

### Figure S12 (related to Figure 1)

Validation of the BRAQUE pipeline analysis.

**A:** Comparison of image segmentation with Cellpose 2.0 and CyBorgh. Three TMA cores (UPN26, 32 and 33) were segmented using Cellpose 2.0 or CyBorgh (part of the BRAQUE pipeline). BRAQUE was run on the .csv data files and the clusters were classified. The mean cell type percentages ± SD are shown. For an explanation of the cell types, see Table S16.

**B:** Spatial and phenotypic discrimination of juxtaposed cells by BRAQUE. Two examples of cells juxtaposed are shown both in the UMAP and in the real space for UPN107. In each panel the clusters of endothelium and stroma (i, gray background), Lyve1+ macrophages and Lyve1+ endothelium (ii, white background) are plotted in contrasting colors, respectively on the UMAP (left) and real space (right). Details are magnified as insets. The markers significance distribution of two representative clusters for each pair are shown below the UMAP maps. Below each real space, a RGB composite of DAPI (blue) and color-coded representative markers for each cell type (red, green) are shown.

**i:** Endothelium (red) and stromal cells (blue) clusters are plotted in the UMAP space and on the real tissue space. The dashed squares are enlarged as inserts. The marker expression ranking of two adjacent clusters shown in the insert, clusters 103 (stromal) and cluster 105 (endothelial) is shown, with relevant markers highlighted. The  triple IF RGB color image shows nuclear DAPI (blue), endothelial vWF (red) and fibroblasts CD248 (green). Note the very close relationship of CD248 fibroblasts with vWF+ endothelium. Scale bar: 500 µm.

**ii:** Lyve1+ Macrophages (red) and Lyve1+ endothelium (blue) clusters are plotted in the UMAP space and on the real tissue space. The dashed squares are enlarged as inserts. The marker expression ranking of two representative clusters, Lyve1+ Macrophages and Lyve1+ endothelium is shown, with relevant markers highlighted. The triple IF RGB color image shows nuclear macrophage PU1 (blue), endothelial Lyve1 (red) and macrophage CD163 (green). Note Lyve1+ and Lyve1- CD163+ macrophages displaying nuclear PU1. Scale bar: 100 µm

**C:** Comparison of cell classification by BRAQUE on AF-subtracted and raw images. 75 images from a TMA core (UPN32) were either autofluorescence-subtracted (noAF) or left untreated (with AF), then both were independently analyzed with BRAQUE and the clusters obtained were classified. The percentage for each cell type is reported.

**D:**Technical artifacts and excluded cells. Clusters classified as technical artifacts (“junk”) are plotted on real space. The scale represents pixels (0.45 µm/pixel). Bottom left: the color-coded clusters plotted on the UMAP space of UPN107. Black dots represent clusters/cells which are allocated in the HDBSCAN -1 cluster. The dotted area is magnified on the right.

**E:** Effect of the antibody panel composition on the detection of cell types by BRAQUE. Two LN (UPN107 and UPN108), stained with 78 markers and segmented, were analyzed with four panels: *i*) BRAQUE with the full panel (78 Abs), *ii*) non redundant, mutually exclusive markers (33 Abs), *iii*) a combination of anti-transcription factor antibodies and 14 key diagnostic markers (TF & friends; 41 Abs) and *iv*) antibodies selected for highest signal-to-noise ratio and cell type identification power (Only cleans; 68 Abs). The percentage of broad cell types in each panel is shown ±SD. Note the larger SD and an increase in junk and unclear clusters with the smaller panels. For details of the panel’s composition see Table S3, S7. Table S7 contains individual cases results with a detailed cell list.

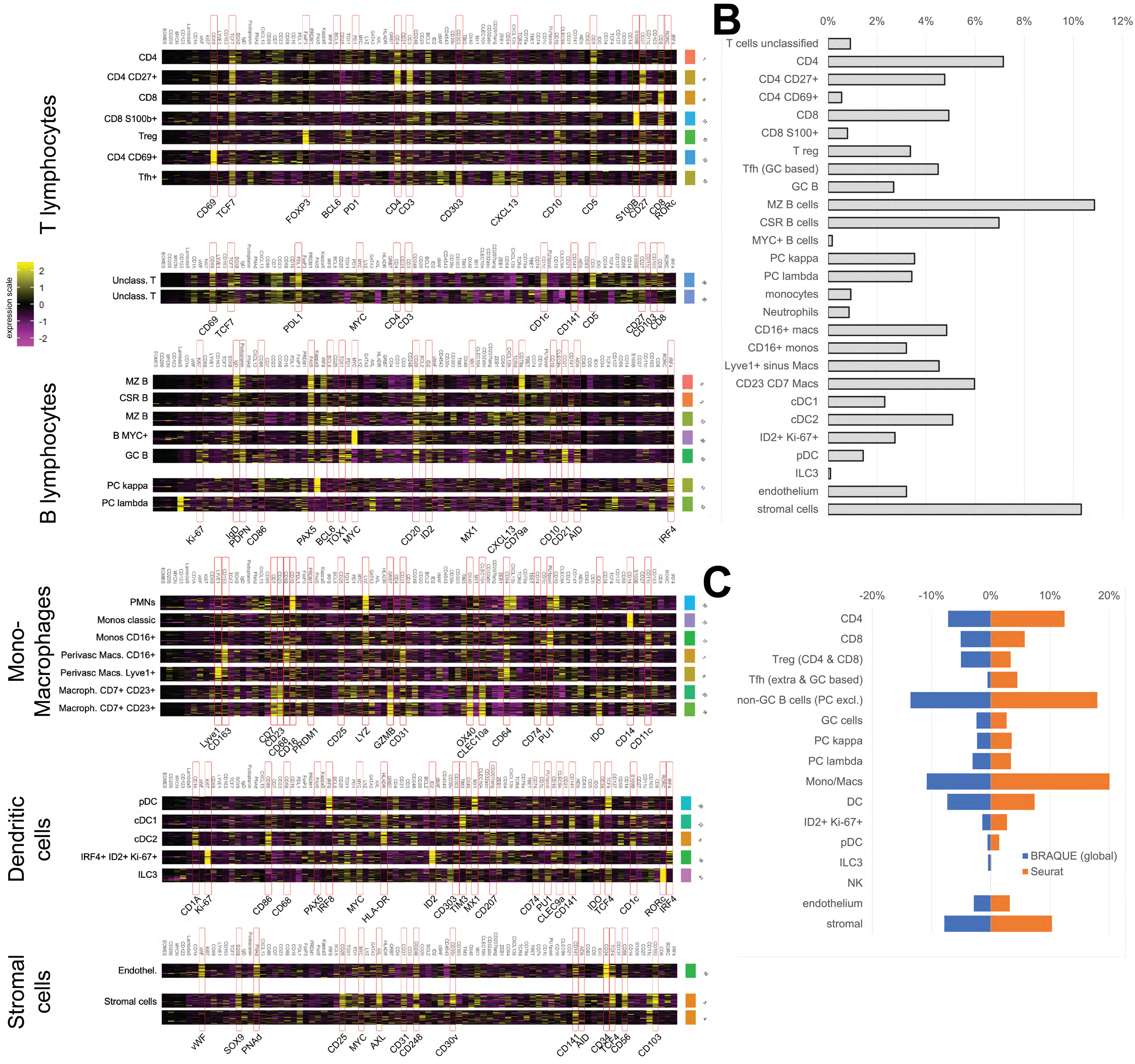


### Figure S13 (related to Figure 1)

Cluster analysis (Seurat) of 37 normal lymph node Tissue Microarray cores.

**A:** 37 TMA cores from 18 subjects stained in multiplex are clustered with Seurat. Individual clusters are grouped by cell type classification. Relevant markers for each group are highlighted (red rectangle and marker name at the bottom). Cluster 45 to 51 have been removed because of insufficient cell numbers each. Unclassifiable clusters 36, 38, 42 and 43 have been removed.

**B:** Cell type composition of 37 TMA cores from 18 subjects are clustered with Seurat. Individual clusters are grouped by cell type classification and the percentage composition is plotted. Note that unclear clusters (199992 cells, 10.7% of the total) have been removed before calculating the percentage of each type. The cell type names are different from the analysis with BRAQUE and are assigned by manual scoring of the heatmaps.

**C:** Comparison of global BRAQUE analysis and Seurat. 37 TMA cores from 18 subjects stained in multiplex are clustered with Seurat and BRAQUE. The cell types from BRAQUE are grouped in broad cell types for the comparison.

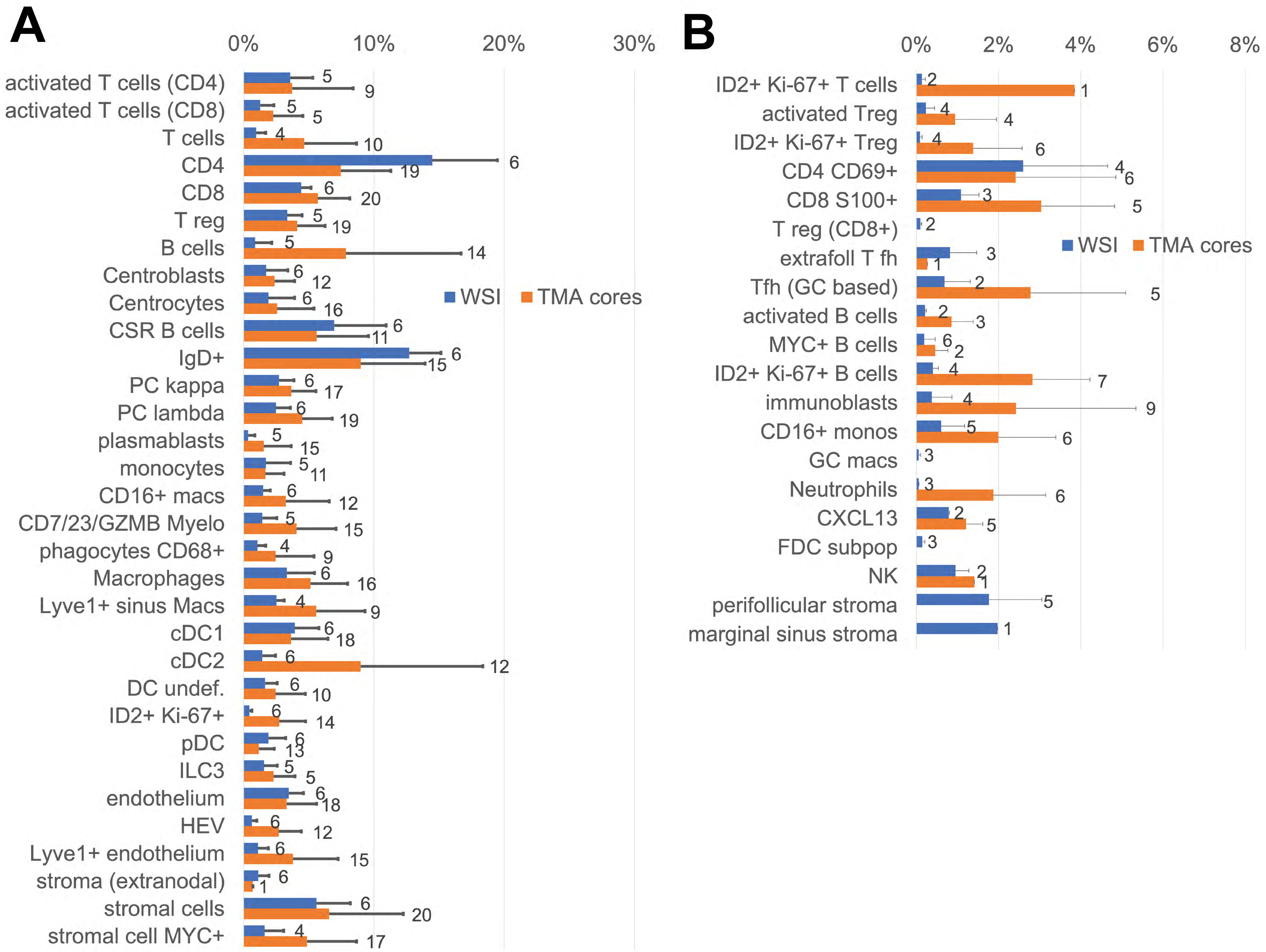


### Figure S14 (related to Figure 1)

Cell type frequency in whole sections (WSI) and TMA cores identified in 27 LN samples (6 whole LN and 21 2mm TMA cores) after BRAQUE$^{global}$ analysis on all cells in each sample, expressed as percentage ± SD. The cell types are separated into **A:** frequent cell types (i.e. ≥ 20 total clusters) and **B:** infrequent cell types (i.e. <20 total clusters). The numbers next to the bars are the number of informative cases for that cell type. Note frequency discrepancies, most notable in **B**, referable to the TMA area sampling selection. The higher frequency of “B cells” (B cells not further identifiable, **A**) in TMAs is due to lack of AID and IgD staining in some TMA cores. See supplemental Tables and Supplemental Data.
